# Inhibition of arterivirus RNA synthesis by cyclophilin inhibitors is counteracted by mutations in replicase transmembrane subunits

**DOI:** 10.1101/587261

**Authors:** Adriaan H. de Wilde, A. Linda Boomaars-van der Zanden, Anja W. M. de Jong, Montserrat Barcéna, Eric J. Snijder, Clara C. Posthuma

## Abstract

Previously, the cyclophilin inhibitors cyclosporin A (CsA) and Alisporivir (ALV) were shown to inhibit the replication of diverse RNA viruses, including arteriviruses and coronaviruses, which both belong to the order *Nidovirales*. Here we aimed to identify arterivirus proteins involved in the mode-of-action of cyclophilin inhibitors and to investigate how these compounds inhibit arterivirus RNA synthesis in the infected cell. Repeated passaging of the arterivirus prototype equine arteritis virus (EAV) in the presence of CsA revealed that reduced drug sensitivity is associated with the emergence of adaptive mutations in nonstructural protein 5 (nsp5), one of the transmembrane subunits of the arterivirus replicase polyprotein. Introduction of singular nsp5 mutations (nsp5 Q21R, Y113H, or A134V) led to a ∼2-fold decrease in sensitivity to CsA treatment, whereas combinations of mutations further increased EAV’s CsA resistance. The detailed experimental characterization of engineered EAV mutants harboring CsA-resistance mutations implicated nsp5 in arterivirus RNA synthesis. Particularly, in an *in vitro* assay, EAV RNA synthesis was far less sensitive to CsA treatment when nsp5 contained the adaptive mutations mentioned above. Interestingly, for increased sensitivity to the closely-related drug ALV CsA-resistant nsp5 mutants required the incorporation of an additional adaptive mutation, which resided in nsp2 (H114R), another transmembrane subunit of the arterivirus replicase. Our study provides the first evidence for the involvement of nsp2 and nsp5 in the mechanism underlying the inhibition of arterivirus replication by cyclophilin inhibitors.

**Importance:** Currently, no approved treatments are available to combat infections with nidoviruses, a group of plus-stranded RNA viruses including important zoonotic and veterinary pathogens. Previously, the cyclophilin inhibitors cyclosporin A (CsA) and Alisporivir (ALV) were shown to inhibit the replication of diverse nidoviruses (both arteriviruses and coronaviruses), and may thus represent a class of pan-nidovirus inhibitors. Here, using the arterivirus prototype equine arteritis virus, we have established that resistance to CsA and ALV treatment is associated with adaptive mutations in two trans-membrane subunits of the viral replication complex, nonstructural proteins 2 and 5. This is the first evidence for the involvement of specific replicase subunits of nidoviruses in the mechanism underlying the inhibition of their replication by cyclophilin inhibitors. Understanding this mechanism of action is of major importance to guide future drug design, both for nidoviruses and other RNA viruses inhibited by these compounds.

## Introduction

Equine arteritis virus (EAV) is a positive-strand RNA (+RNA) virus that belongs to the arterivirus family in the order *Nidovirales*, a steadily expanding clade of positive-stranded RNA (+RNA) viruses that also includes the roni-, mesoni-, and coronavirus families (https://talk.ictvonline.org/ictv-reports/ictv_9th_report/positive-sense-rna-viruses-2011/w/posrna_viruses/219/nidovirales). The latter group includes the zoonotic severe acute respiratory syndrome coronavirus (SARS-CoV) and Middle East respiratory syndrome (MERS)-CoV, which can cause lethal respiratory infections in humans (reviewed by (Cui et al., 2018; de Wit et al., 2016)). Furthermore, the nidovirus order includes several important veterinary pathogens, such as avian, porcine, and bovine coronaviruses, and the arterivirus porcine reproductive and respiratory syndrome virus (PRRSV). The latter continues to cause major economic losses in the swine industry worldwide (Holtkamp et al., 2013). Despite extensive efforts during recent years, currently no approved treatments to combat nidovirus infections are available.

Equine arteritis virus (EAV) is the prototype of the arterivirus family and has a genome size of 12.7 kilobases (kb) (Snijder et al., 2013). About three-quarters of its polycistronic genome is occupied by the viral replicase gene, which is expressed as two polyproteins (pp), pp1a and the C-terminally extended pp1ab, with the synthesis of the latter depending on a programmed ribosomal frameshift that can occur just upstream of the ORF1a stop codon. The post-translational maturation of pp1a and pp1ab (Fig. 1A) involves extensive proteolytic cleavage (into at least 13 non-structural proteins (nsps)) by viral proteases residing in nsp1 (PLP1), nsp2 (PLP2), and nsp4 (M^pro^). In the cytoplasm of arterivirus-infected cells, most of the viral nsps and viral RNA synthesis are associated with membranous ‘replication organelles’ (ROs) that typically include paired membranes and double-membrane vesicles (DMVs; (Knoops et al., 2012; Zhang et al., 2018), and reviewed by (van der Hoeven et al., 2016)). Co-expression of the nsp2 and nsp3 transmembrane subunits of the EAV replicase induces the formation of similar double-membrane structures (Snijder et al., 2001). Recently, a role in modulating RO membrane curvature and DMV formation was proposed for the third transmembrane replicase subunit, nsp5 (van der Hoeven et al., 2016). The core of the arteriviral RNA-synthesizing machinery, sometimes also referred to as replication and transcription complex (RTC), is formed by ORF1b-encoded nsps like nsp9 and nsp10, which include the RNA-dependent RNA-polymerase (RdRp) and helicase (Hel) domains, respectively. In addition, host factors appear to be required to promote efficient viral RNA synthesis (van Hemert et al., 2008). The RTC directs viral genome replication as well as the synthesis of a nested set of subgenomic (sg) mRNAs that is used to express the genes encoding the viral structural proteins, which are located in the 3’-proximal quarter of the genome (reviewed in (Snijder et al., 2013)).

**Fig. 1.**
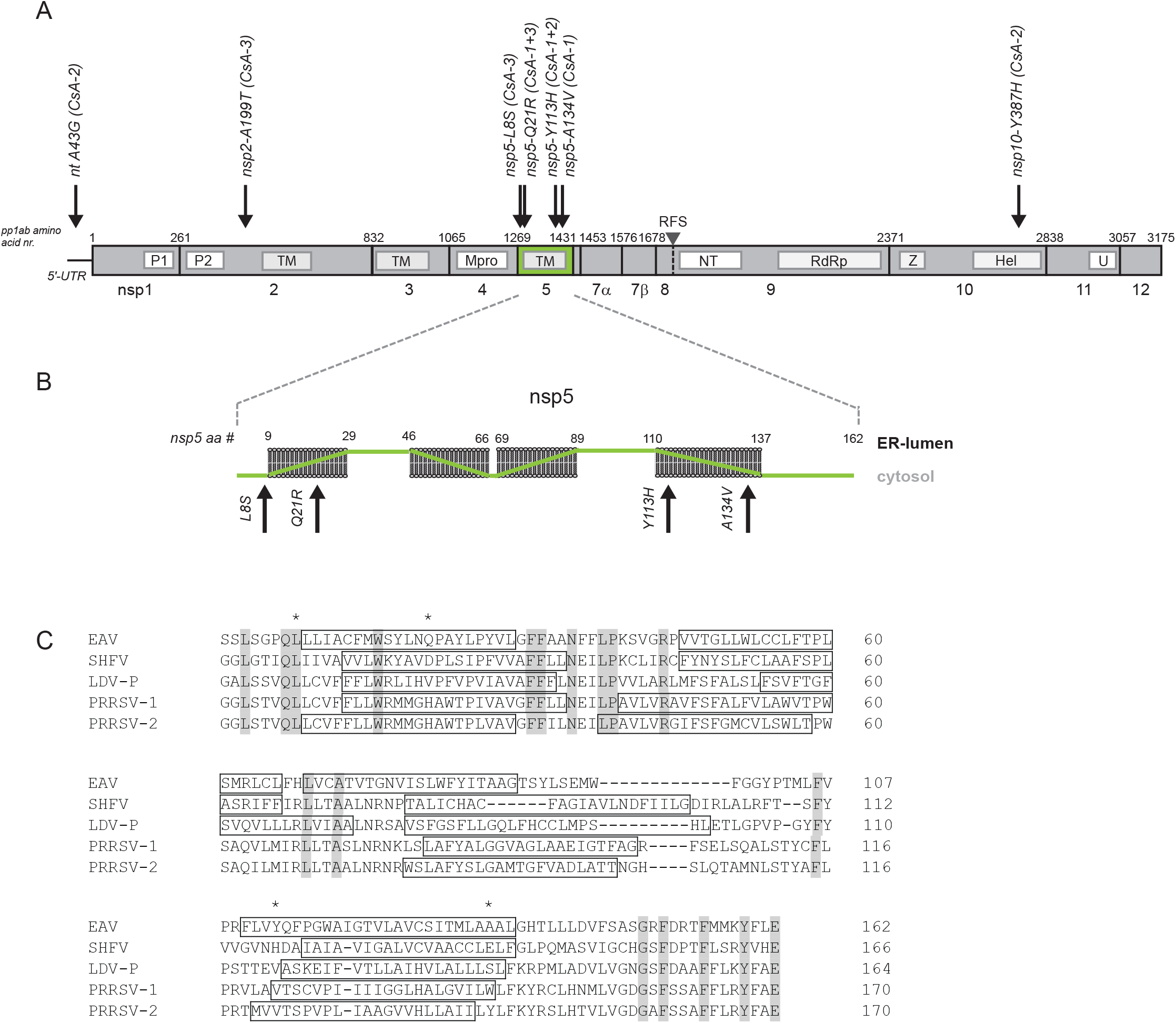
Positions of CsA resistance-associated mutations in the EAV 5’UTR and ORF1ab. (A) Map of CsA resistance-associated mutations identified in the consensus sequence of lineages CsA-1, -2, and -3 after seven passages. Depicted are the 5’-UTR and ORF1ab, with replicase polyprotein 1ab amino acid numbers, cleavage sites and nsp cleavage products indicated. The seven mutations identified in the sequences of the CsA-resistant viruses are depicted with black arrows, and the amino acid position/change in the respective nsp is indicated, as well as the lineage in which the mutation was identified (see also Table 2). Nsp5 is highlighted in green. P1 and P2: papain-like proteinases in nsp1 and nsp2, resp.; TM: transmembrane domains; Mpro: main proteinase; NT: nidovirus RdRp- associated nucleotidyltransferase (NiRAN); RdRp: RNA-dependent RNA polymerase; Z: Zinc-binding domain; Hel: helicase; U: endoribonuclease. (B) Membrane topology of EAV nsp5 as predicted using the TMHMM topology prediction method within the Geneious software package. The position/change of the four nsp5 mutations identified in lineages CsA-1, -2, and -3 is indicated with a black arrow. (C) Multiple sequence alignment of nsp5 from selected arteriviruses, performed by Clustal Omega (Chojnacki et al., 2017). Fully conserved residues are indicated in grey, predicted transmembrane helices (TMHMM topology prediction from Geneious software package) are boxed and adaptive mutations are marked with an asterisk. EAV, equine arteritis virus (GenBank accession number DQ846750); SHFV, simian hemorrhagic fever virus (AF180391); PRRSV-1, porcine reproductive and respiratory syndrome virus, European genotype (GU737264.2); PRRSV-2, porcine reproductive and respiratory syndrome virus, North American genotype (JX138233); LDV, lactate-dehydrogenase-elevating virus (U15146).

Multiple laboratories, including our own, have shown that in cell culture-based infection models low-micromolar concentrations of the FDA-approved cyclophilin inhibitor cyclosporin A inhibit the replication of a wide variety of nidoviruses, including the arteriviruses EAV and PRRSV (de Wilde et al., 2013a) as well as SARS-CoV, MERS-CoV, and other CoVs (de Wilde et al., 2013b; de Wilde et al., 2011; Pfefferle et al., 2011; Tanaka et al., 2013)). CsA inhibits members of the cyclophilin (Cyp) family, which are peptidyl-proline isomerases (PPIases) that act as chaperones and facilitate protein folding and function (reviewed in (Dunyak and Gestwicki, 2016; Wang and Heitman, 2005)). However, CsA also has immune-suppressive properties (Schreiber and Crabtree, 1992) that would constitute a highly undesirable side-effect in the context of antiviral therapy. Therefore, numerous non-immunosuppressive Cyp inhibitors have been developed. One of these, Alisporivir (ALV) has been identified as a potent inhibitor of hepatitis C virus (HCV) and other RNA viruses (reviewed in (de Wilde et al., 2018a; Naoumov, 2014)). Previously, we showed that also nidovirus replication is sensitive to ALV treatment, using drug concentrations comparable to the EC_50_ previously determined for CsA (de Wilde et al., 2017).

The most ubiquitously expressed member of the Cyp family is the cytosolic CypA, which has been proven to be an essential host factor in the replication of *e.g*. hepatitis C virus (HCV), human immunodeficiency virus 1 (HIV-1), West Nile virus (reviewed in (Frausto et al., 2013)), as well as the coronaviruses HCoV-NL63 and HCoV-229E (Carbajo-Lozoya et al., 2014; von Brunn et al., 2015). However, other coronaviruses seem to be much less affected by depletion of CypA in the infected cell (de Wilde et al., 2011); (de Wilde et al., 2013a; de Wilde et al., 2018b). EAV replication, on the other hand, strongly depends on CypA expression, as it was nearly abolished in CypA-knockdown or – knockout cell cultures (de Wilde et al., 2013a; de Wilde et al., 2018b).

Our present study aimed to investigate the molecular details of the inhibition of arterivirus replication by cyclophilin inhibitors. To this end, we first passaged EAV in the presence of inhibitory CsA concentrations, to induce resistance to the compound. Subsequently, adaptive mutations that markedly decreased the sensitivity of EAV replication to CsA treatment were found to map to nsp5, one of the transmembrane subunits of the arterivirus replicase. We could show that CsA treatment interferes with EAV RNA synthesis, and that the CsA-resistance mutations in nsp5 render replication, and in particular RNA synthesis, less sensitive to CsA treatment. Furthermore, we show that resistance to the closely related Cyp inhibitor ALV requires an additional adaptive mutation in nsp2, a second transmembrane subunit of the arterivirus replicase that is essential for both polyprotein processing and RO formation. Our studies implicate both nsp2 and nsp5 in the mechanism underlying the sensitivity of arterivirus replication to cyclophilin inhibitor treatment. Moreover, they suggest that the role of cyclophilins as arterivirus host factors is specifically linked to viral RNA synthesis, possibly through interactions with key players in RO formation.

## Material and methods

### Cell culture, EAV infection, and virus titration

BHK-21 cells (Nedialkova et al., 2010), Huh7 cells (de Wilde et al., 2013b), and 293T cells (van Kasteren et al., 2012) were cultured as described previously. A cell culture-adapted derivative of the EAV Bucyrus isolate (Bryans et al., 1957) was used to infect BHK-21 and Huh7 cell monolayers at 37°C as described previously (de Vries et al., 1992; Nedialkova et al., 2010); (van der Hoeven et al., 2016). EAV titers in cell culture supernatants were determined by plaque assay on BHK-21 cells (Nedialkova et al., 2010). Huh7 cells lacking CypA expression (Huh7-CypA^KO^ cells), generated by transfection with a pLentiCRISPR v2 plasmid and a CypA-specific guide RNA, were described previously (de Wilde et al., 2018b).

### Isolation of drug-resistant EAV mutants

To obtain CsA- or ALV-resistant mutants, EAV was passaged (MOI 0.005) in Huh7 cells in the presence of increasing compound concentrations (4 to 20 µM for CsA, 4 to 75 µM for ALV). For CsA, passaging was repeated independently three times, resulting in EAV lineages CsA-1, CsA-2, and CsA-3. For ALV, two independent ALV-resistant lineages (EAV ALV-1 and ALV-2) were obtained. A control lineage (EAVwt), passaged in the absence of CsA or ALV, was generated in parallel to screen for adaptive mutations unrelated to drug treatment, which might arise during cell culture passaging of EAV. CsA or ALV concentrations were increased in the following passage at the moment that in drug-treated, EAV-infected cells, virus-induced CPE appeared at the same moment as in the control-passaged EAV lineage, indicating that virus replication is not affected by that particular drug concentration. Drug-specific toxicity was monitored in cells that were left uninfected. Resistance to CsA or ALV was monitored using the cell-based antiviral screening assay described below.

### Cell culture-based assay for inhibition of EAV infection

Huh7 cells were seeded in transparent 96- well plates at a density of 10^4^ cells per well. After overnight incubation at 37°C, 100 µl of 1.5x concentrated compound dilution and 50 µl of EAV inoculum in DMEM–2% FCS were added to each well. The MOI used was 0.05 and the final compound concentrations in the medium that were tested ranged from 0.4 to 50 µM. DMSO (0.25%, CsA) or ethanol (0.25%, ALV) were used as solvent control. At 3 days post infection (p.i.), differences in cell viability due to virus-induced cytopathic effect (CPE) or compound-specific side effects were analyzed using the CellTiter 96 AQ_ueous_ Non-Radioactive Cell Proliferation Assay (Promega), as described previously (de Wilde et al., 2013b). The cytotoxic effects of treatment with compound only were monitored in parallel in plates containing mock-infected cells. The half-maximal effective concentration (EC_50_) and the compound-specific toxicity (50% cytotoxic concentration; CC_50_) were calculated with GraphPad Prism 6 software, using the non-linear regression model.

### Identification of adaptive mutations in the EAV genome

To isolate viral RNA, Huh7 cells (10-cm^2^ dish) were infected with wild-type (wt) EAV, wtEAV P7, EAV CsA lineages, or EAV ALV lineages (MOI of 0.1). Cells were incubated for 16 h at 37°C, lysed in 1 ml of TriPure (Sigma Aldrich), and intracellular RNA was isolated according to the manufacturer’s instructions. Reverse transcription (RT) was performed with the ThermoScript™ RT-PCR System (ThermoFisher Scientific) according to the manufacturer’s instructions, using random hexamers (Promega) and 10 µM of primer 5’-CGCCGTTTTTTTTTTTTTTTTTTTTTTTTT-3’. Subsequently, the complete EAV genome was amplified as seven PCR products and sequenced (primer sequences used in RT, PCR and sequencing are available upon request).

### EAV reverse genetics

Resistance-associated adaptive mutations (listed in Table 2) were introduced into shuttle plasmids by standard site-directed PCR mutagenesis. After sequence verification, restriction fragments containing the desired mutations were transferred to full-length EAV cDNA clone pEAN900, a derivative of pEAV030 (van Dinten et al., 1997) containing a few translationally silent mutations to engineer restriction sites. Virus derived from clone pEAN900 has a wt phenotype, as confirmed by a side-by-side comparison with pEAV030-derived virus in growth curve experiments and plaque assays (data not shown). *In vitro* transcribed RNA derived from wt or mutant pEAN900 was electroporated into BHK-21 cells using Amaxa Nucleofector technology (Lonza), as described previously (Beerens and Snijder, 2007). Recombinant virus was harvested upon complete CPE, typically between 48 and 72 h post transfection (p.t.). The presence of the engineered mutations was confirmed by performing one additional virus passage and sequencing RT-PCR products obtained after isolation and RT-PCR amplification of intracellular viral RNA.

### Electron microscopy

Huh-7 cells were infected with rEAV^wt^ or rEAV^QYA^, or were mock-infected, and either treated with 8 µM CsA at 1 h p.i. or left untreated. Cells were fixed at 11 (no CsA) or 14 h p.i. (8 µM CsA) in 1.5% glutaraldehyde in 0.1 M cacodylate buffer (pH 7.4) for 1 h. Subsequently, samples were stained during 1-h incubation steps, first with 1% OsO_4_ in 0.1 M cacodylate buffer and then with 1% uranyl acetate. Samples were then dehydrated in a graded ethanol series (70% to 100% ethanol) and embedded in Epon LX-112 resin (Ladd Research) that was polymerized at 60°C. Sections of 100 nm thickness were collected on carbon-coated pioloform EM grids and post-stained with lead citrate and a saturated aqueous solution of uranyl acetate. Images were collected on a Tecnai12 TWIN electron microscope (Thermo Fisher Scientific (formerly FEI)) operating at 120 kV using a OneView 4k high frame-rate camera (Gatan).

### Isolation of EAV RTC-containing replication organelles and *in vitro* RNA synthesis assays

Membrane-associated EAV RTCs were isolated from BHK-21 cells and *in vitro* RNA synthesis assays (IVRAs) were performed essentially as described previously (van Hemert et al., 2008). In short, approximately 5 × 10^7^ EAV-infected BHK-21 cells (MOI 5) were harvested at 7.5 h p.i. and lysed using a ball-bearing homogenizer (Isobiotek) with 16 µm clearance. Lysates were centrifuged for 10 min at 1000 × g to obtain a post nuclear supernatant (PNS). A standard IVRA contained PNS (the equivalent of 6 × 10^4^ infected cells) and a 5x concentrated CsA stock (final concentration 4 to 32 µM) or RTC dilution buffer (control) and was performed in the presence of [α-^32^P]CTP that was incorporated in the newly synthesized EAV RNA. Assays were incubated at 30°C for 100 min and terminated by the addition of 60 µl of 5% LiDS-LET-ProtK. After a 15-min incubation at 42°C, the ^32^P-labeled reaction products were isolated and separated in a 1.5% denaturing agarose gel. Agarose gels were dried and reaction products were visualized by phophorimaging using a Typhoon 9410 imager (GE Healthcare). Incorporation of label was quantified using ImageQuant TL software.

### Analysis of viral RNA synthesis

Metabolic labeling of viral RNA with [^3^H]uridine was performed essentially as described previously (Knoops et al., 2010). Briefly, 4 × 10^5^ BHK-21 cells in 4-cm^2^ wells were infected with rEAV^wt^ or rEAV^QYA^(MOI 5). At 5.5 h p.i., DMEM containing 8% FCS, 5 µg/ml dactinomycin (Sigma-Aldrich), and 8 µM CsA or 0.02% DMSO (solvent control) was given. At 6.5 h p.i., 100 µCi of [^3^H]uridine was added to 300 µl of culture medium. Cells were lysed at 7.5 h p.i. and intracellular ^3^H-labeled RNAs were isolated (Tripure), separated by denaturing agarose gel electrophoresis, and visualized by fluorography. The total incorporation of ^3^H label was determined using liquid scintillation counting. For hybridization analysis of intracellular RNA from EAV-infected cells, RNA was first separated by denaturing RNA agarose gel electrophoresis. Subsequently, the gel was dried and hybridization was performed using a ^32^P-labeled oligonucleotide probe (5’-TTGGTTCCTGGGTGGCTAATAACTACTT-3’) recognizing the 3’end of the EAV genome, as described previously (van Hemert et al., 2008).

## Results

### Selection of CsA-resistant EAV mutants

Previously, we showed that low-micromolar CsA concentrations can effectively block EAV and PRRSV replication in cultured cells (de Wilde et al., 2013a). We now aimed to investigate the mechanism-of-action by which arterivirus infection is inhibited in the presence of this cyclophilin inhibitor. To this end, wt EAV (strain Bucyrus; MOI 0.005) was serially passaged in Huh7 cells in the presence of increasing concentrations of CsA (4 to 20 µM). Three independent virus lineages were generated (named EAV CsA-1 to -3), which exhibited clearly decreasing sensitivity to CsA (Fig. 2A). By passage 7 (P7), all three lineages replicated in the presence of 20 µM CsA (data not shown), a dose that is approximately 10-fold higher than the EC_50_ value observed when using wtEAV (Fig. 2B and (de Wilde et al., 2013a)). The CsA sensitivity of the three lineages was measured in an assay that is based on (the inhibition of) EAV-induced CPE in Huh7 cells. Following infection with an MOI of 0.05 in the presence of 0.4 to 50 µM CsA, cell viability was assayed at 3 days p.i. CsA doses above 12.5 µM appeared to be cytotoxic, as the viability of mock-infected CsA-treated cells dropped below 85% compared to Huh7 cells not treated with CsA (Fig. 2B; light grey line). The CPE induced by wtEAV infection was inhibited by CsA with an EC_50_ value of 2.4 µM, whereas treatment with 3.1 µM CsA fully blocked the development of virus-induced CPE. This was in line with our previous results in EAV-infected BHK-21 cells, in which treatment with 4 µM CsA resulted in a 4-log reduction of viral progeny (de Wilde et al., 2013a). Whereas seven passages in the absence of CsA did not alter the drug sensitivity of the wtEAV control (EC_50_ 2.1 µM), P7 virus of the CsA-treated EAV lineages CsA-1 to -3 had become five-fold less sensitive to the compound (Fig. 2B and Table 1). Of note, additional passaging (up to passage 9) did not decrease CsA sensitivity any further.

**Table 1:** CsA sensitivity of serially passaged and engineered CsA-resistant viruses.

|  | EC<sub>50</sub> | Fold resistance** | Remarks |
| --- | --- | --- | --- |
| **wild-type EAV** |  |  |  |
| wtEAV | 2.4 ± 0.5 | 1.0 |  |
| wtEAV P7 | 2.1 ± 0.2 | 1.0 |  |
| rEAVwt | 1.6 ± 0.1 | 1.0 |  |
| **EAV CsA-1** |  |  |  |
| Passaged virus P7 | ~10.9* | 5.1 |  |
| rEAV nsp5<sup>Q21R</sup> | 4.4 ± 0.2 | 2.9 |  |
| rEAV nsp5<sup>Y113H</sup> | 4.4 ± 0.3 | 2.8 |  |
| rEAV nsp5<sup>A134V</sup> | 4.5 ± 0.3 | 2.9 |  |
| rEAV nsp5<sup>Y113H+A134V</sup> | 9.1 ± 0.6 | 5.9 | Acquisition of second site reversion nsp5 Q21R or Q21H |
| rEAV nsp5<sup>Q21R+Y113H+A134V</sup> (rEAV<sup>QYA</sup>) | 7.2 ± 0.7 | 4.7 |  |
| **EAV CsA-2** |  |  |  |
| Passaged virus P7 | ~11.0* | 5.2 |  |
| rEAV<sup>A43G</sup> | 1.8 ± 0.1 | 1.2 |  |
| rEAV nsp5<sup>Y113H</sup> | 4.4 ± 0.3 | 2.8 |  |
| rEAV nsp10<sup>Y387H</sup> | 1.6 ± 0.1 | 1.1 |  |
| rEAV<sup>A43G</sup>+nsp5<sup>Y113H</sup>+nsp10<sup>Y387H</sup> | 4.6 ± 0.3 | 3.0 |  |
| **EAV CsA-3** |  |  |  |
| Passaged virus P7 | ~10.2* | 4.8 |  |
| rEAV nsp2<sup>A199T</sup> | 1.7 ± 0.1 | 1.1 |  |
| rEAV nsp5<sup>L8S</sup> | 2.2 ± 0.1 | 1.4 | Reversion to wt nsp5-L8 |
| rEAV nsp5<sup>Q21R</sup> | 4.4 ± 0.2 | 2.9 |  |
| rEAV nsp5<sup>L8S+Q21R</sup> | 9.1 ± 1.4 | 5.9 | Reversion to wt nsp5-L8 |
| rEAV nsp2<sup>A199T</sup>+nsp5<sup>L8S+Q21R</sup> | 8.7 ± 0.5 | 5.6 | Reversion to wt nsp5-L8 |

\* The EC<sub>50</sub> is an estimated value, since a reliable calculation could not be performed due to the low viability at the maximal inhibitory ALV concentration.
\*\* Fold resistance of the P7 viruses relative to wtEAV; for the engineered rEAV ALV mutants fold resistance was calculated relative to the rEAV<sup>wt</sup> value.

**Table 2:**
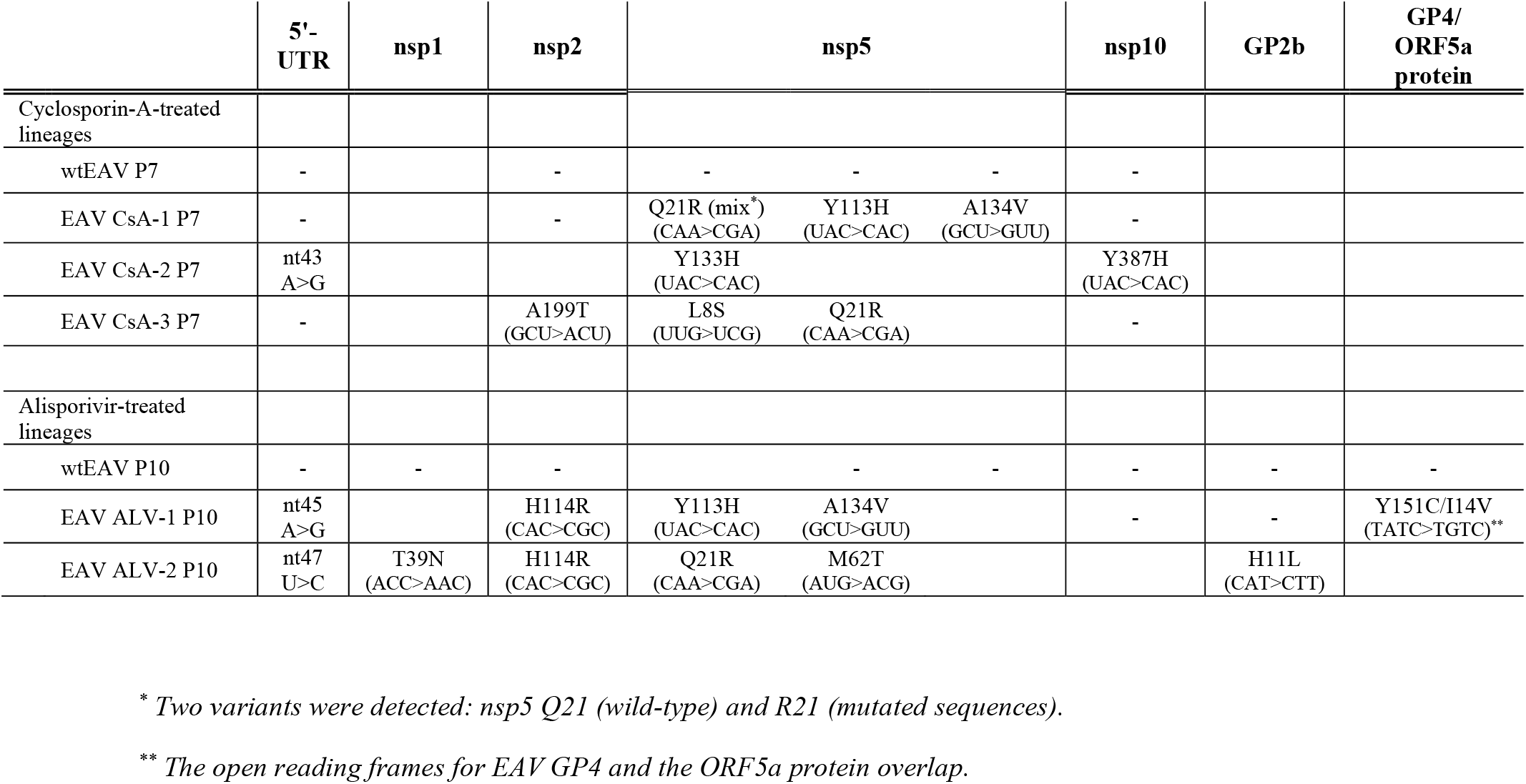
Mutations identified after serial passaging of EAV in the presence of increasing concentrations of CsA or ALV

**Fig. 2.**
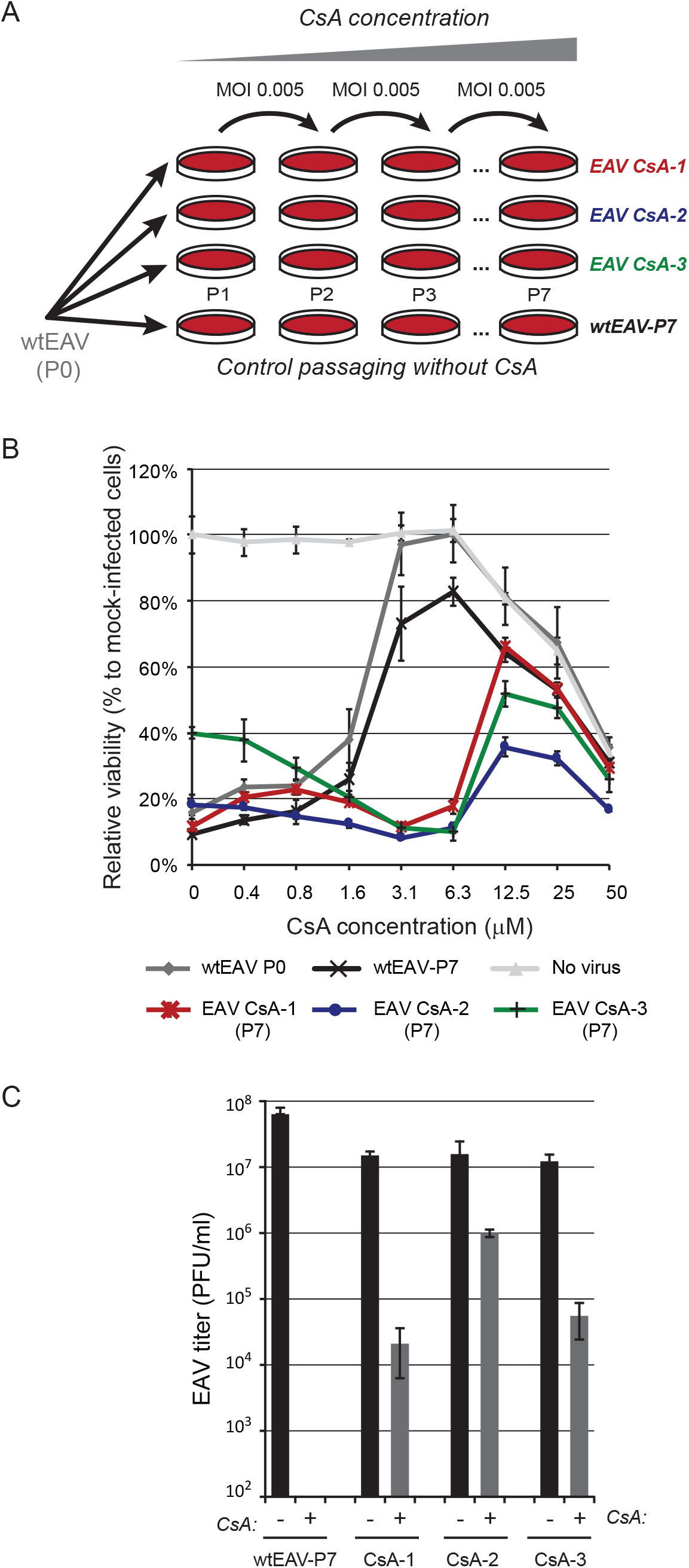
Selection of CsA-resistant EAV mutants. (A) Passaging scheme for wild-type (wt) EAV (strain Bucyrus) in Huh7 cells (MOI 0.005) to generate three independent CsA-resistant lineages (EAV CsA-1 in red; CsA-2 in blue; and CsA-3 in green). Passaging was performed in the presence of increasing CsA concentrations, ranging from 4 µM during passage 1 (P1) to 20 µM during P7. As a control wtEAV (black) was passaged in the absence of CsA to assess the acquisition of mutations unrelated to CsA resistance. (B) After P7, EAV CsA-1 (red), CsA-2 (blue) and CsA-3 (green) were tested for CsA resistance, while including the original (P0, grey) and passaged (P7, black) wtEAV as controls. Huh7 cells in 96-well plates were infected at MOI 0.05 in the presence of 0 to 50 µM CsA. Cells were incubated for three days and cell viability was monitored using a commercial assay. Furthermore, the cytotoxicity of CsA treatment only was monitored in parallel in mock-infected Huh7 cells (light grey). Graphs show the results (averages and standard deviations [SD]) of a representative experiment performed in quadruplicate. All experiments were repeated at least twice. (C) Huh7 cells were infected with EAV CsA-1, −2, or −3 or with wtEAV-P7 (MOI 0.01). At 1 h p.i., the inoculum was replaced by medium containing either 0.02% DMSO (bars labeled -; solvent control) or 4 µM CsA (bars labeled +). Medium was collected at 32 h p.i. and EAV progeny titers (in PFU/ml) were determined by plaque assay.

Next, we confirmed the reduced CsA sensitivity of CsA-1 to -3 P7 virus by infecting (MOI 0.01) Huh7 or BHK-21 cells in the presence of 4 µM CsA, a dose that fully inhibits wtEAV replication. Virus production was measured at 48 h p.i., which confirmed that wtEAV-P7 infection in the presence of 4 µM CsA indeed did not yield detectable virus progeny (Fig. 2C). At the same CsA concentration, titers of above 10^4^ PFU/ml (EAV CsA-1 and CsA-3) or even around 10^6^ PFU/ml (EAV CsA-2) were reached when infection was done with the P7 viruses from the three CsA-resistant lineages. Similar results were obtained in BHK-21 cells, albeit that the yields of all three lineages were equally reduced compared to wtEAV-P7 in this cell line (data not shown).

### Most CsA resistance-associated mutations map to EAV nsp5

In order to identify CsA resistance-associated mutations, the consensus genome sequence of the three P7 CsA-resistant EAV samples was determined following RT-PCR amplification (Table 2 and Fig. 1A). The EAV CsA-1 consensus sequence contained three mutations, all mapping to the nsp5-coding region of ORF1a and resulting in three amino acids substitutions: Q21R (mixed with the wt sequence), Y113H, and A134V. Remarkably, the Y113H mutation in nsp5 was also encoded by the ORF1a consensus sequence of the EAV CsA-2 population, but now accompanied by a mutation in the 5’ untranslated region (5’-UTR) (A43G) and one in ORF1b, specifying a mutation in the nsp10 helicase domain (Y387H). Lineage CsA-3 contained mutations leading to substitutions in nsp2 (A199T) and again nsp5 (L8S and Q21R). None of the consensus sequences contained any synonymous mutations that had been fixed during passaging. Likewise, no mutations were discovered in the consensus sequence of the wtEAV-P7 control, implying that the mutations identified in the CsA-1 to -3 P7 viruses potentially are linked to the established reduced sensitivity to CsA.

To assess the importance of the identified mutations for CsA sensitivity, each mutation was individually reverse engineered into the EAV genome, using full-length cDNA clone pEAN900. Recombinant mutant viruses were launched and their CsA sensitivity was assayed in Huh7 cells using the CPE-based assay outlined above (see Fig. 1b). Interestingly, three of the nsp5 mutations (Q21R, Y113H, and A134V) decreased the CsA sensitivity of EAV replication (EC_50_ values of ∼4.4 µM vs. 1.6 µM for the wt control (rEAV^wt^)). No change in sensitivity was observed for any of the other mutations (see Table 1 and sFig. 1A-H). Of note, the nsp5^L8S^ mutation reverted back to the wild-type residue (data not shown), which likely explains the marginal decrease in CsA sensitivity observed in this assay (sFig. 1C).

The increase in CsA resistance observed for the recombinant mutant viruses with singular mutations appeared to be ∼3-fold smaller than the approximately 5-fold increase observed for the P7 CsA-1 to -3 viruses, which each carried multiple substitutions. Therefore, we also engineered mutant viruses with genome sequences identical to the consensus sequence of these three CsA-resistant virus lineages. For lineage CsA-1, which contained a mix of Q (wt) and R (mutant) at residue 21 of nsp5, both variants were engineered and tested. Compared to the viruses carrying singular mutations, mutant rEAV nsp5^Y113H+A134V^ displayed a decreased sensitivity to CsA, which now approached that of the EAV CsA-1 lineage (Fig. 3A and Table 1). When the Q21R mutation was added, the CsA resistance diminished slightly compared to the Y113H+A134V double mutant (Fig. 3B and Table 1; EC_50_ of 9.1 vs. 7.2 µM). However, upon passaging and resequencing (data not shown), the double mutant appeared to be genetically instable, which may also explain the mixed sequence found at nsp5 codon 21 in the lineage CsA-1 consensus sequence. For lineage CsA-2, combining the three mutations (in 5’- UTR, nsp5, and nsp10) into the same genome did not further improve CsA resistance compared to rEAV nsp5^Y113H^ (Fig. 3C). This lack of a synergistic effect indicates that the mutations in 5’-UTR and nsp10 do not contribute substantially to the level of CsA resistance of CsA-2. In the case of EAV CsA-3, combining nsp5^L8S^, but not nsp2^A199T^, with nsp5^Q21R^ yielded a ∼2-fold further decrease in CsA sensitivity (EC_50_ value of 9.1 µM; compare sFigs. 1B, E and Fig. 3D and see Table 1) suggesting that the nsp2^A199T^ mutation does not contribute to CsA resistance.

**Fig. 3.**
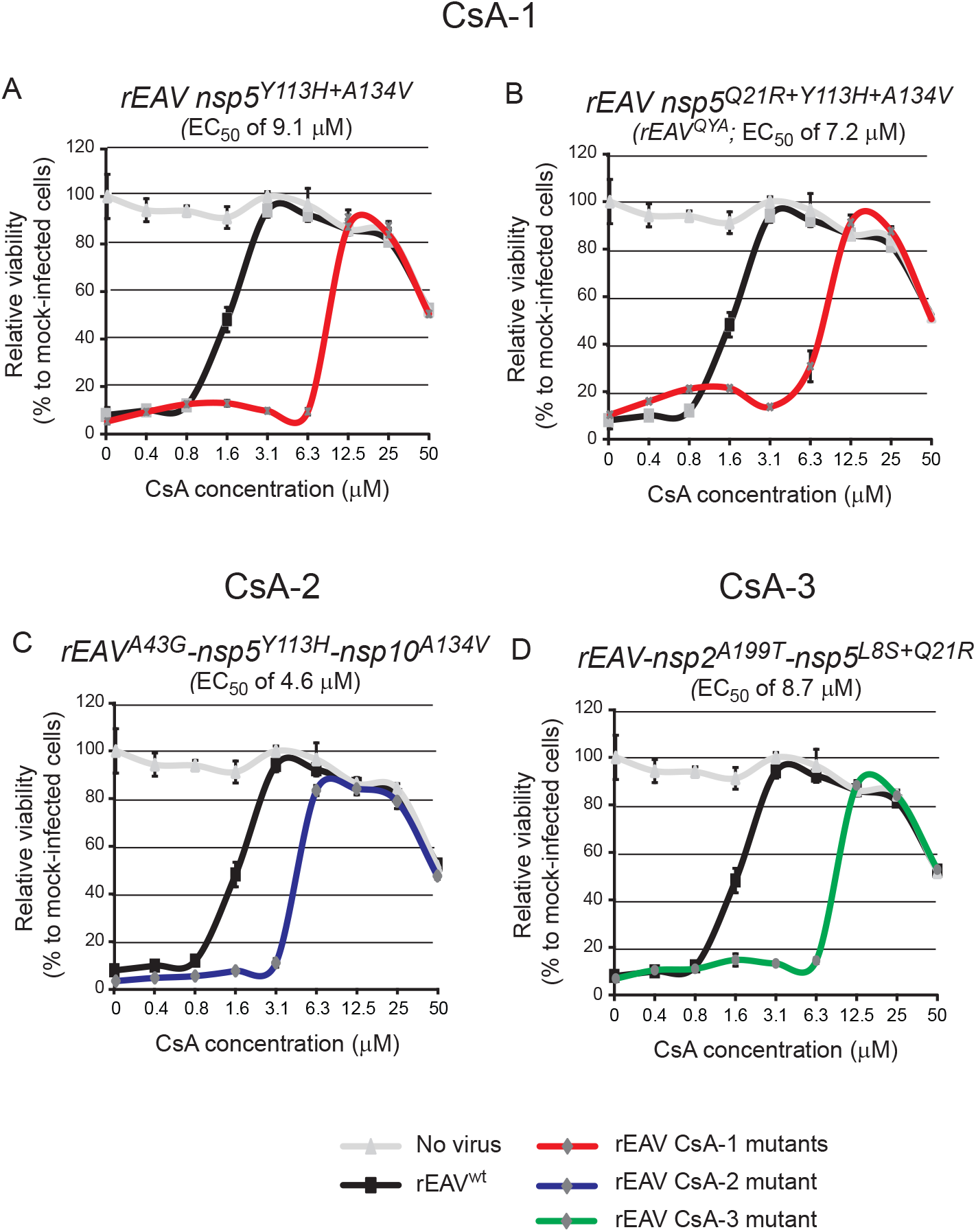
CsA resistance analysis of engineered rEAV mutants reflecting the consensus sequences of CsA-1, -2, and -3. Huh7 cells in 96-well plates were infected (MOI 0.05) with rEAV^wt^ (control; black line) or mutants reflecting the consensus sequences of CsA-1, -2, and -3 (red lines in panels A and B, blue line in C and green line in D). Cells were incubated for three days in the presence of 0 to 50 µM CsA, and cell viability was monitored using a commercial assay. The cytotoxicity of CsA treatment only was monitored in parallel in mock-infected Huh7 cells (light grey line). Above each panel, the EC_50_ value of CsA inhibition is indicated (see also Table 1). Graphs show the results (average and SD) of a representative experiment that was performed in quadruplicate. All experiments were repeated at least twice.

Overall, the above results clearly implicated various EAV nsp5 mutations in reduced CsA sensitivity. Whereas combining nsp5 mutations increased CsA resistance compared to singular nsp5 mutants, the substitutions in 5’-UTR, nsp2, and nsp10 did not contribute detectably to CsA resistance. CsA-1 mutant rEAV nsp5^Q21R+Y113H+A134V^ was selected for follow-up experiments, as it was genetically stable upon passaging (data not shown) and included the single nsp5 mutation (Y113H) that was key to the CsA resistance of lineage CsA-2. For simplicity, this mutant will be referred to as rEAV^QYA^ from this point forward.

### rEAV^QYA^ replicates in the presence of CsA

The CsA resistance of rEAV^QYA^ was analyzed in more detail by infecting Huh7 cells in the presence or absence of 4 µM CsA. Upon high-MOI inoculation (MOI 3), the replication kinetics of rEAV^wt^ and rEAV^QYA^ in the absence of CsA were very similar (Fig. 4A; solid lines). Progeny was released from rEAV^wt^- and rEAV^QYA^-infected cells from 12 h p.i. onwards, with rEAV^QYA^ titers being only slightly reduced compared to those for rEAV^wt^. In the presence of 4 µM CsA, rEAV^wt^ was essentially unable to replicate (Fig. 4A; black dashed line) while rEAV^QYA^ was able to produce infectious progeny, albeit with a delay compared to the CsA-free infection (Fig. 4A; red dashed line). Subsequently, the CsA resistance of rEAV^QYA^ was further confirmed in a low-MOI infection experiment (MOI 0.01) in Huh7 cells (Fig. 4B): a titer of about 10^4^ PFU/ml was obtained for the mutant in the presence of 4 µM CsA, while rEAV^wt^ infectious progeny was not detected under the same conditions.

**Fig. 4.**
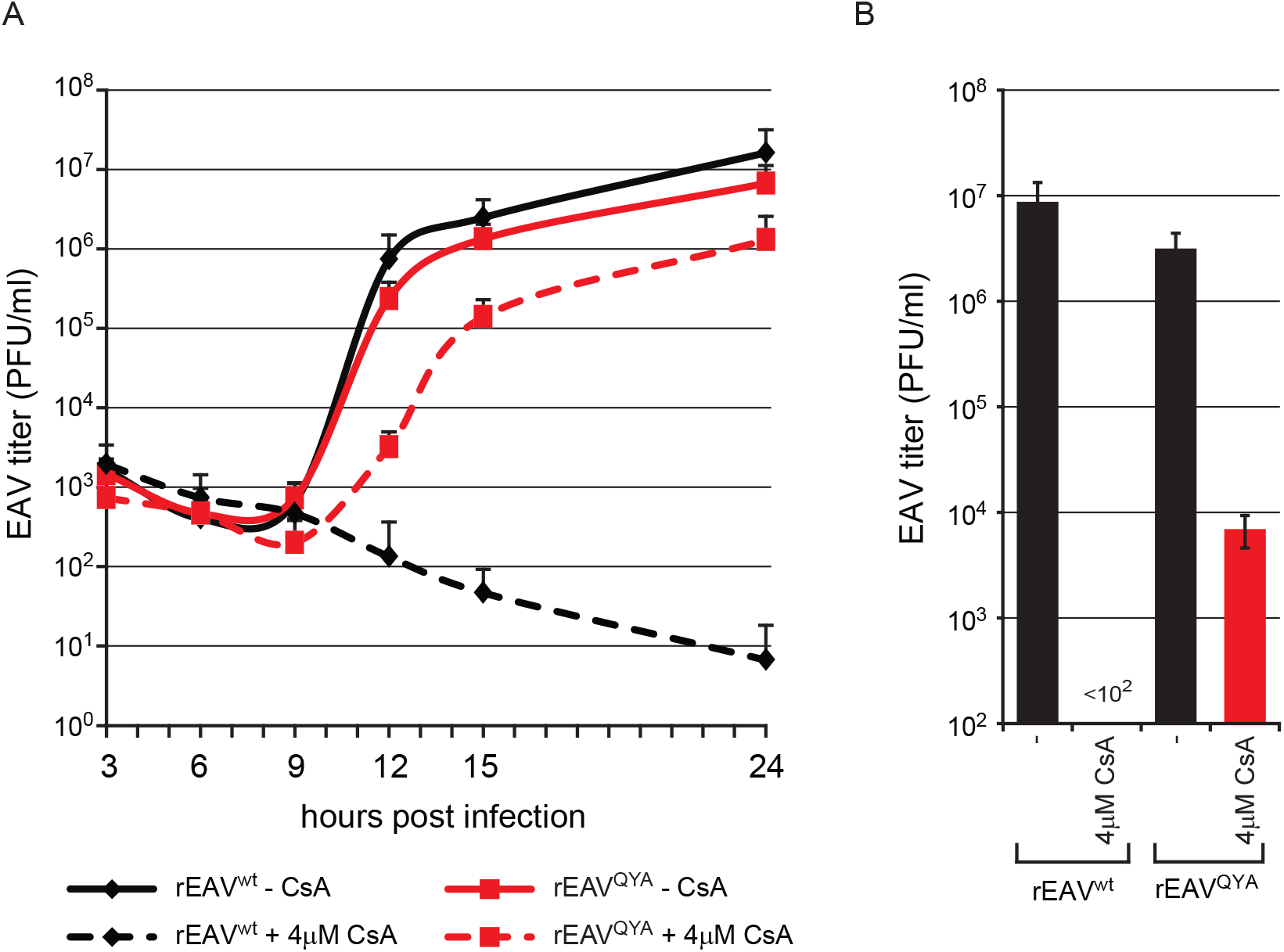
Virus yields of rEAV^wt^ and rEAV^QYA^ infection in the absence or presence of CsA. (A) Growth kinetics of rEAV^wt^ (black lines) and rEAV^QYA^ (red lines) in Huh7 cells (MOI 5). Infections were performed in the absence (solid lines) or presence of 4µM CsA (dotted lines). EAV yields at the indicated time points were determined by plaque assay (averages and SD given, n=3). (B) Virus yields of rEAV^wt^ and rEAV^QYA^ (MOI 0.01) in Huh7 cells at 32 h p.i. in the absence or presence of 4µM CsA. EAV yields were determined by plaque assay (average and SD given, n=3).

### Neither CsA treatment nor nsp5 mutations affect the morphology of EAV-induced replication organelles

It was previously reported that Cyp inhibitor treatment affects the morphology of the ‘membranous web’, the RO that is formed upon HCV infection and with which viral RNA synthesis is associated (Chatterji et al., 2015; Madan et al., 2014). It was striking that the EAV mutations that promote CsA-resistance all map to the nsp5 transmembrane protein, which has been suggested to play a regulatory role in the formation of arterivirus ROs (van der Hoeven et al., 2016), of which the peculiar DMVs are a key feature (Pedersen 1999, Knoops 2012). We therefore used electron microscopy to investigate whether CsA treatment or the mutations in nsp5 affect EAV-induced DMV formation. Analysis of Huh7 cells (11 h p.i.) infected with either rEAV^wt^ (Fig. 5B) or rEAV^QYA^ (Fig. 5C) did not reveal obvious differences in DMV amounts, size, or ultrastructure. As replication of rEAV^QYA^ in the presence of CsA was delayed compared to the replication in untreated cells, we analyzed cells infected with this mutant at 14 h p.i (Fig. 4). In the presence of CsA, no DMVs were observed for rEAV^wt^ (Fig. 5E), while the morphology and size of the DMVs in rEAV^QYA^-infected cells were similar to the structures observed upon infection of untreated cells infected with rEAV^wt^- or rEAV^QYA^ (Fig. 5F). Notably, severe compound-induced dilation of ER membranes was observed, a phenomenon that was independent of EAV infection as it was also observed in CsA-treated mock-infected Huh7 cells (compare Figs. 5A-C and 5D-F). Taken together, these observations strongly suggest that CsA treatment interferes with a very early stage of arterivirus infection, *i.e*. prior to or during RO formation. Apparently, adaptive mutations in nsp5 can compensate for the detrimental effects of CsA treatment, which may either target one of the viral players directly or affect a host factor involved in arterivirus replication, like cyclophilin A (see above).

**Fig. 5.**
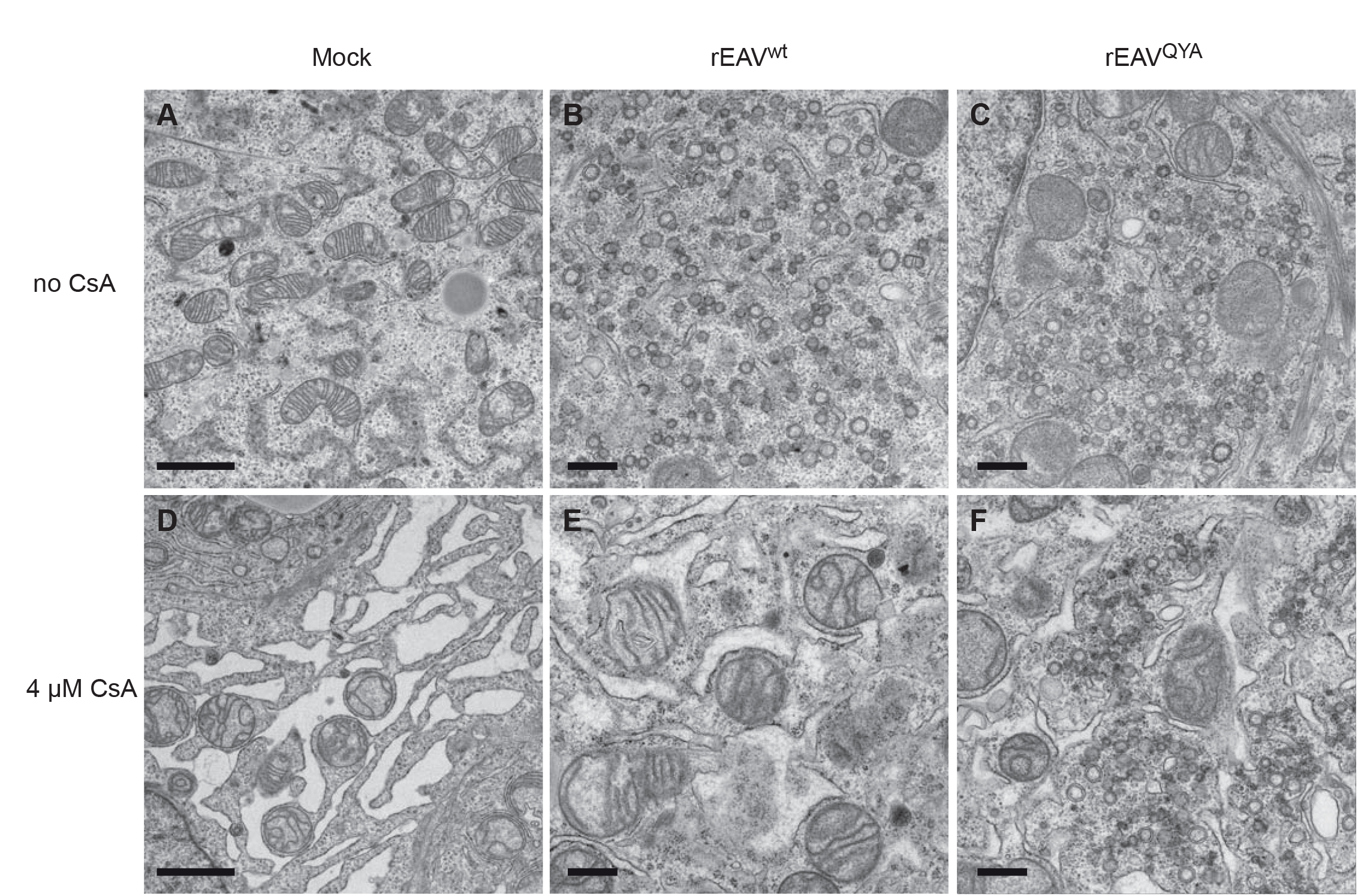
Adaptive mutations in EAV nsp5 do not affect the morphology of virus-induced double membrane vesicles. Huh7 cells were infected with rEAV^wt^ (panels B and E) or rEAV^QYA^ (panels C and F) with MOI 5, or were mock-infected (panels A en D). After an incubation of 11 h (untreated, panels A-C) or 14 h (CsA-treated, panels D-F), cells were fixed and processed for transmission electron microscopy. DMV formation is very similar upon infection with rEAV^wt^ and rEAV^QYA^, while CsA treatment induces dilated ER, both in mock and in infected cells. Scale bars: 2 µm (left panels), 500 nm (middle and right panels).

### rEAV^QYA^ replication continues to depend on CypA

We previously showed that EAV replication was reduced in cells depleted for CypA by siRNA treatment (de Wilde et al., 2013a; de Wilde et al., 2018b) and strongly affected in a CypA knock-out cell line (Huh7-CypA^KO^) produced using CRISPR- Cas9 technology (de Wilde et al., 2018b). These latter cells were used to investigate whether the replication of rEAV^QYA^ still depended on CypA expression. We infected both the Huh7-CypA^KO^ cells and the parental Huh7 cells with wt and mutant virus (MOI 0.01) and compared progeny titers at 32 h p.i. At this time point, rEAV^wt^ yields were almost 4-log reduced in the knockout cells (Fig. 6). Replication of rEAV^QYA^ was inhibited to the same extent, indicating that – despite its reduced CsA sensitivity – replication of this mutant continues to depend on the presence of CypA (Fig. 6).

**Fig. 6.**
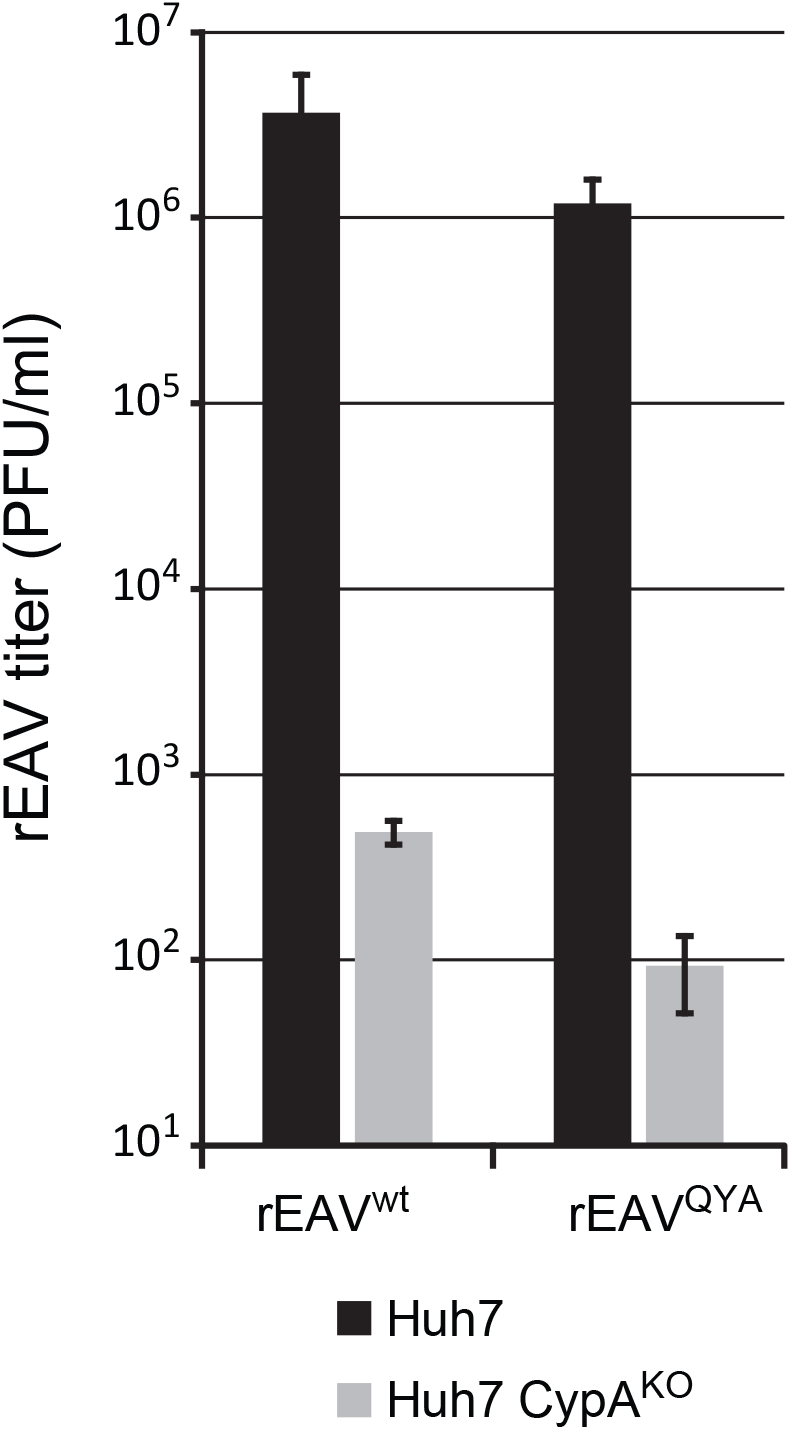
rEAV^QYA^ does not replicate in the absence of CypA. (A) Virus yields of rEAV^wt^ (left) and rEAV^QYA^ (right; MOI 0.01) at 32 h p.i. of parental Huh7 cells (black bars) and Huh7 CypA^KO^ cells (grey bars). EAV yields were determined by plaque assay (averages and SD, n=3).

### Nsp5 mutations reduce the inhibitory effect of CsA on EAV RNA synthesis

Previously, we showed that EAV RNA synthesis is effectively inhibited by CsA (de Wilde et al., 2013a). This led us to analyze whether the nsp5^QYA^ mutations reduce the inhibitory effect of CsA on EAV RNA synthesis. For this purpose, BHK-21 cells were infected with rEAV^wt^ or rEAV^QYA^ and viral RNA synthesis was assayed by metabolic labelling with [^3^H]uridine from 6.5 to 7.5 h p.i. (Fig. 7A, B). Cells were also treated with dactinomycin to block cellular transcription. [^3^H]-uridine incorporation into viral RNA was quantified by isolation of intracellular RNA and liquid scintillation counting (Fig. 7A). Labeled viral RNA was also visualized by agarose gel electrophoresis and fluorography (Fig. 7B). This revealed that RNA synthesis by the mutant virus was approximately 40% of wt virus activity, which is in line with the observed delayed growth kinetics and reduced titers of the mutant. However, in the presence of 8 µM of CsA (which was added 1 h prior to the pulse labeling), rEAV^wt^ RNA synthesis was completely blocked, while rEAV^QYA^ RNA synthesis was hardly affected, as evident from both the overall quantification of ^3^H-labeled RNA and the RNA analysis using agarose gel electrophoresis (Fig. 7A-B).

**Fig. 7.**
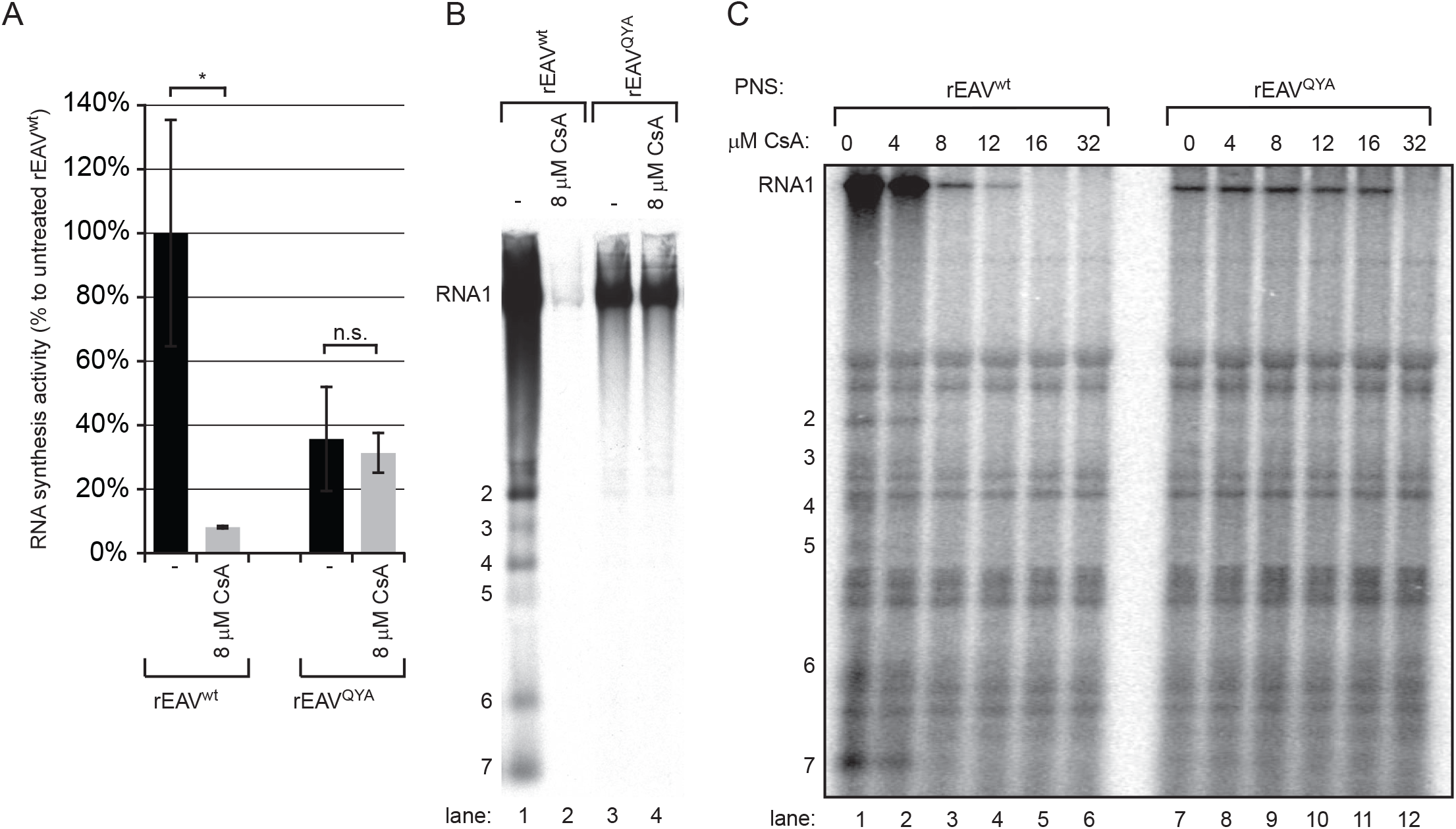
rEAV^QYA^ RNA synthesis is impaired but less sensitive to CsA. (A, B) Viral RNA synthesis in BHK-21 cells infected with rEAV^wt^ or rEAV^QYA^ was metabolically labeled between 6.5 and 7.5 h p.i. using [^3^H]uridine. This was done in the presence or absence of 8 µM CsA and dactinomycin. Total intracellular RNA was isolated at 7.5 h p.i. (A) The total incorporation of ^3^H label was quantified by liquid scintillation counting. (B) ^3^H-labelled EAV RNA was separated in a denaturing formaldehyde agarose gel and visualized by fluorography. (C) ROs were semi-purified from lysates of rEAV^wt^- or rEAV^QYA^-infected BHK-21 cells (MOI 5) at 7.5 h p.i. and were used in an *in vitro* RNA synthesis assay (IVRA) in which [^32^P]CTP was incorporated into viral RNA products. Reactions were performed in the presence of increasing concentrations of CsA (indicated above the lanes) and were terminated after 100 min. Labeled RNA products were isolated, separated in a denaturing formaldehyde agarose gel, and visualized by phosphor imaging. The positions of the genomic RNA (RNA1) and subgenomic RNAs (positions 2 to 7) are indicated on the left side of the gel.

Next, we assayed the RNA-synthesizing activity of semi-purified ROs from rEAV^wt^- and rEAV^QYA^-infected cells using a previously developed *in vitro* RNA synthesis assay (IVRA) (van Hemert et al., 2008). The incorporation of [^32^P]CTP into viral RNA was analyzed in the presence of various CsA concentrations. In the absence of the compound, *in vitro* synthesis of rEAV^wt^ genomic and sg RNAs was observed (Fig. 7C, lane 1), which was clearly reduced when the assay was performed in the presence of ≥8 µM of CsA (lanes 3 to 6). In line with the [^3^H]-uridine metabolic labeling experiment, the RNA-synthesizing complexes from rEAV^QYA^-infected cells were insensitive to treatment with up to 16µM of CsA (lanes 7 to 12), thus directly linking the effect of the adaptive nsp5 mutations to the overall activity of the arterivirus RTC.

Of note, the results from both the metabolic labeling with [^3^H]uridine and the IVRA experiments suggested that – compared to genome replication – sg RNA synthesis is more impaired for the rEAV^QYA^ mutant (Fig 7B, lane 1 and 3; Fig. 7C, lane 1 and 7). However, when rEAV^wt^ and rEAV^QYA^ RNA (isolated from infected BHK-21 cells at 7.5 h p.i.) was subjected to direct hybridization analysis using a [^32^P]-labeled probe that recognizes all EAV mRNAs, an overall decrease in the amount of mutant viral RNAs was visible, but no clear difference in the ratio between the individual sg RNAs and genomic RNA (sFig. 2). Therefore, we concluded that the absence of the sg RNAs in some of the panels presented in Fig. 7 must have been due to the limited sensitivity of these particular assays rather than a specific impairment of sg RNA synthesis.

### Resistance to the non-immunosuppressive CsA-analog Alisporivir requires a combination of mutations in EAV nsp5 and nsp2

Previously, we established that EAV replication can also be inhibited by the non-immunosuppressive CsA analog Debio-064 (de Wilde et al., 2013a). More recently, we reported the inhibition of coronavirus replication in cell culture by the related CsA analog Alisporivir (ALV) (de Wilde et al., 2017), a drug that was explored as a host-directed antiviral treatment option for chronic HCV infection (Naoumov, 2014). ALV lacks the immunosuppressive properties of CsA, while retaining a high affinity for cyclophilins. We now established that ALV is able to block also the replication of wtEAV with an EC_50_ value of 4.5 ± 0.2 µM, an efficiency that is comparable to that observed for CsA (Table 3).

**Table 3:** ALV sensitivity of serially passaged and engineered ALV-resistant viruses.

|  | EC<sub>50</sub> | Fold resistance** |
| --- | --- | --- |
| **wild-type EAV** |  |  |
| wtEAV | 4.5 ± 0.2 | 1.0 |
| rEAV<sup>wt</sup> | 2.3 ± 0.3 | 1.0 |
| **EAV ALV-1** |  |  |
| Passaged virus P10 | 33.8 ± 5.8 | 7.4 |
| rEAV nsp2<sup>H114R</sup> | 4.6 ± 0.1 | 2.0 |
| rEAV nsp5<sup>A134V</sup> | 4.3 ± 0.1 | 1.9 |
| rEAV nsp2<sup>H114R</sup> + nsp5<sup>A134V</sup> | 8.9 ± 0.3 | 3.9 |
| rEAV nsp2<sup>H114R</sup> + nsp5<sup>A134V</sup> + GP4<sup>Y151C</sup> | 9.7 ± 0.5 | 4.2 |
| **EAV ALV-2** |  |  |
| Passaged virus P10 | ~12.7 * | 2.8 |
| rEAV nsp2<sup>H114R</sup> | 4.6 ± 0.1 | 2.0 |
| rEAV nsp5<sup>Q21R+M62T</sup> | 3.5 ± 0.2 | 1.9 |
| rEAV nsp2<sup>H114R</sup> + nsp5<sup>Q21R+M62T</sup> | 14.3 ± 1.0 | 6.2 |
| rEAV nsp2<sup>H114R</sup> + nsp5<sup>Q21R+M62T</sup> + GP2<sup>H11L</sup> | 13.2 ± 5.5 | 5.8 |
| rEAV<sup>QYA</sup> + nsp2<sup>H114R</sup> | ~20.0 * | 8.7 |

\* The EC<sub>50</sub> is an estimated value, since a reliable calculation could not be performed due to the low viability at the maximal inhibitory ALV concentration.
\*\* Fold resistance of both P10 viruses relative to wtEAV; for the engineered rEAV ALV mutants fold resistance was calculated relative to the rEAV<sup>wt</sup> value.

To explore the development of ALV resistance in EAV, two independent passaging experiments in Huh7 cells were performed (EAV ALV-1 and ALV-2) in the presence of increasing ALV concentrations (from 4 µM in passage 1 to 75 µM in passage 10), similar to the approach used to generate CsA-resistant viruses (Fig. 2). By passage 10, a CPE-based assay revealed that, compared to the wtEAV-P10, the sensitivity to ALV was reduced by 7.4- and 2.8-fold for EAV lineages ALV-1 and ALV-2, respectively (Fig. 8A and Table 3). Interestingly, the P10 samples of EAV lineages ALV- 1 and −2 were ∼4-fold less sensitive to CsA as well (Fig. 8B). Analysis of the ALV-1 and −2 P10 consensus sequences revealed that each population contained multiple mutations (Table 2). These included singular mutations in the 5’-UTR, in ORF1a (nsp1 region), and in the structural protein-coding region. Strikingly, the P10 viruses from both lineages also carried two substitutions in nsp5. The two nsp5 mutations in ALV-1 (Y113H and A134V) had previously been identified in lineage CsA-1 (see above), whereas ALV-2 also contained a mutation present in CsA-1, Q21R. Strikingly, both ALV-resistant lineages contained the same mutation (H114R) within the papainlike protease (PLP2) domain of nsp2. We reverse engineered the ALV-1 and −2 mutations into the viral genome, and assayed ALV sensitivity of the mutant viruses. Singular nsp2 and nsp5 mutants displayed a reduced ALV sensitivity, but only the nsp5 mutants were also less sensitive to CsA (Fig. 9, Table 3). Subsequently, combining nsp5 and nsp2 mutations had a synergistic effect and confirmed the importance of the common nsp2^H114R^ substitution for ALV resistance. Mutations in the 5’UTR and nsp1 did not further reduce the sensitivity to ALV (data not shown), nor did the mutations in the structural protein-coding region contribute to ALV resistance (Fig. 9; Table 3).

**Fig. 8.**
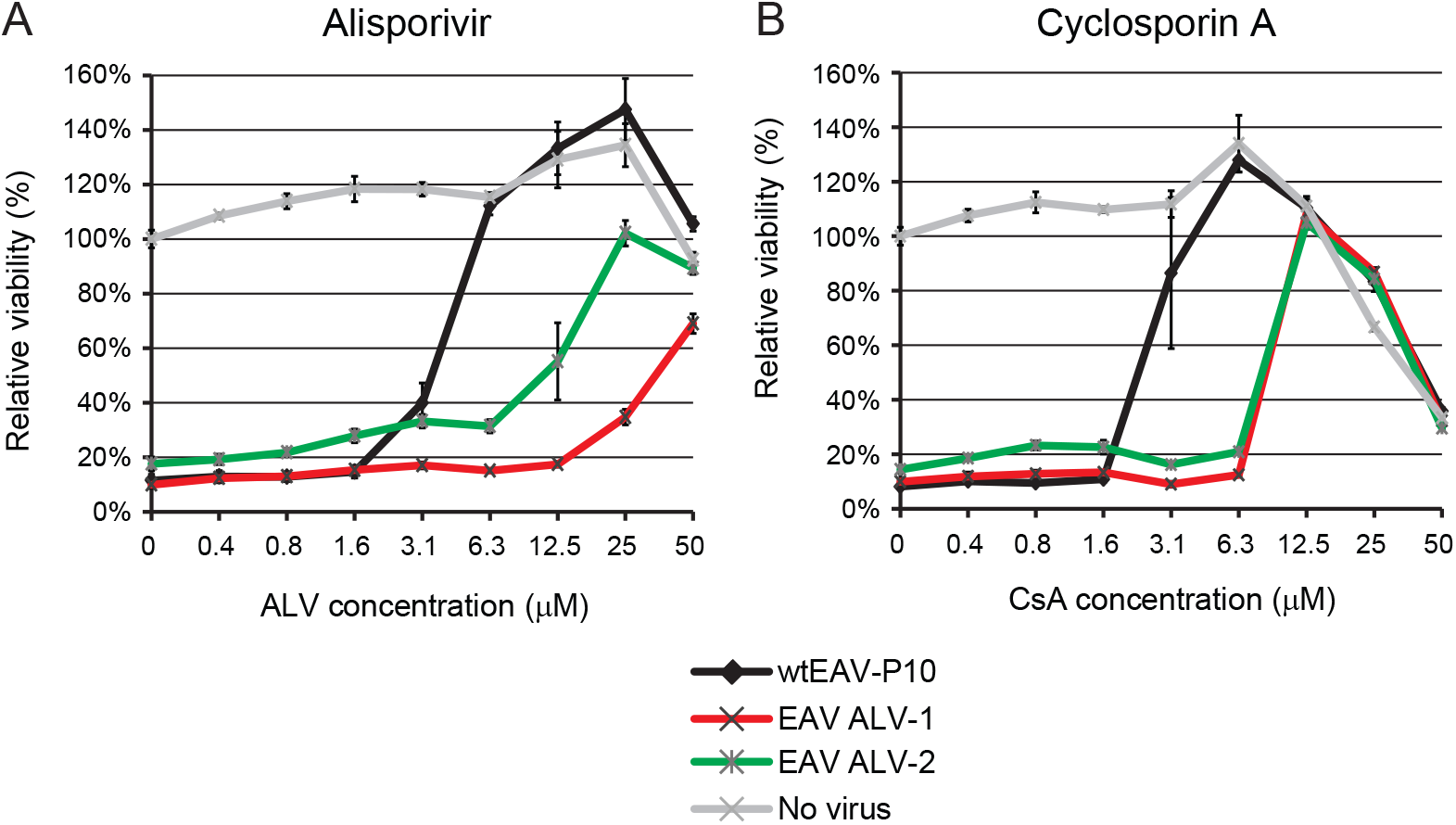
Passaging in the presence of ALV reduces EAV sensitivity towards ALV and CsA. Virus was passaged of increasing ALV concentrations, ranging from 4 µM during passage 1 (P1) to 75 µM during P10. P10 harvests of EAV lineages ALV-1 and ALV-2 were tested for resistance to ALV (A) and CsA (B) treatment. Huh7 cells in 96-well plates were infected with MOI 0.05 in the presence of 0 to 50 µM ALV or CsA. Cells were incubated for three days, and cell viability was monitored using a commercial assay. In addition, the cytotoxicity of CsA treatment was monitored in parallel in mock-infected Huh7 cells. Graphs show the results (average and standard deviations [SD]) of a representative experiment that was performed in quadruplicate. All experiments were repeated at least twice.

**Fig. 9.**
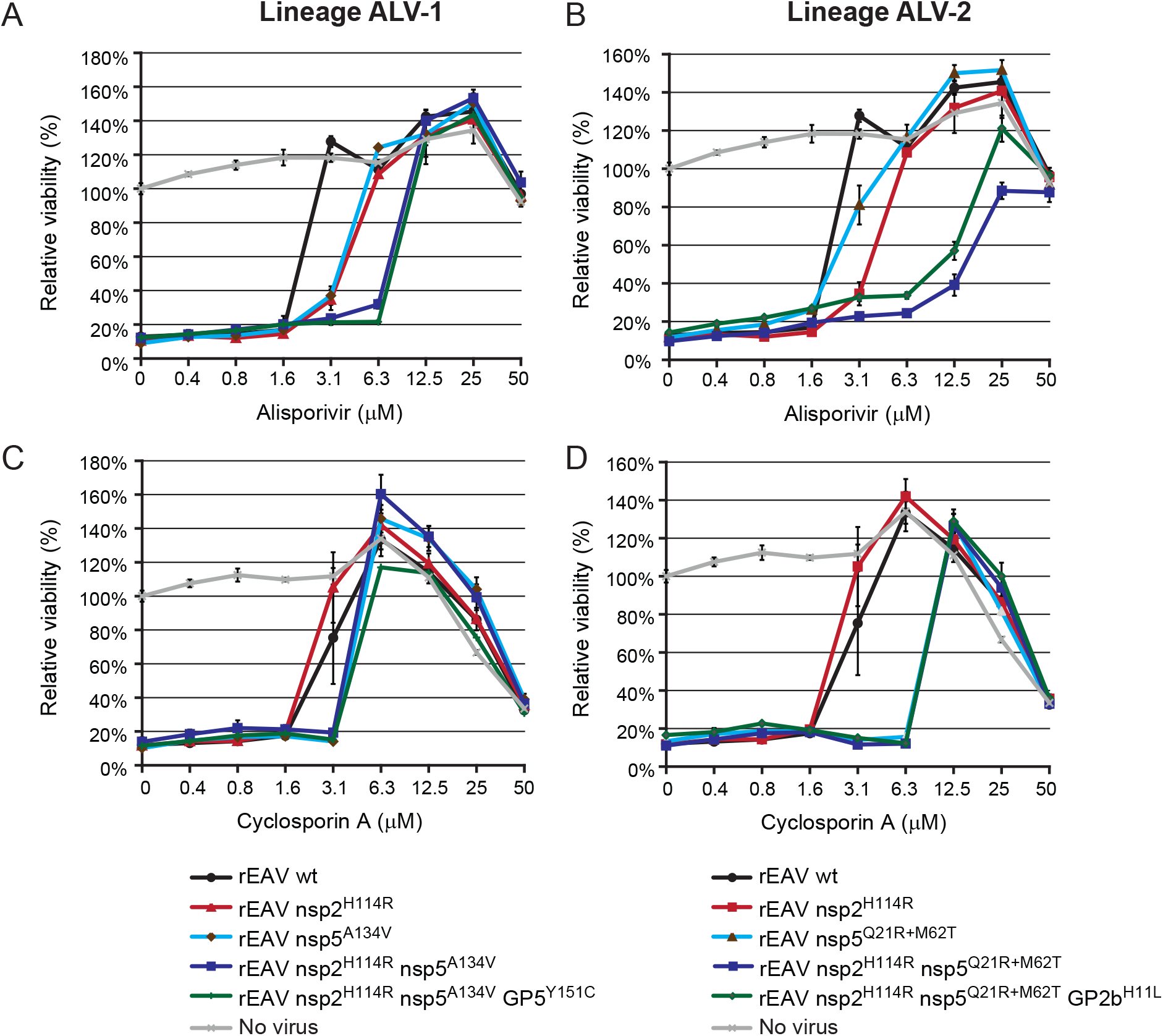
Analysis of ALV and CsA sensitivity of engineered rEAV ALV-resistant mutants. Huh7 cells in 96-well plates were infected with rEAV^wt^ (control) and rEAV ALV-resistant mutants (MOI 0.05), grouped per ALV lineage, in the presence of 0 to 50 µM ALV (A, B) or CsA (C, D). Cells were incubated for three days and cell viability was monitored using a commercial assay. The cytotoxicity of ALV or CsA treatment only was monitored in parallel in mock-infected Huh7 cells (light grey line). See legend for color coding of the rEAV mutants. Graphs show the results (average and SD) of a representative experiment that was performed in quadruplicate. All experiments were repeated at least twice.

Finally, to further corroborate the importance of nsp2^H114R^ for ALV resistance, this mutation was transferred to the rEAV^QYA^ triple mutant already carrying the nsp5 mutations identified in our studies into CsA resistance (see above). As anticipated, while the triple mutant was marginally resistant to ALV (most likely due to the presence of the nsp5^A134V^ mutation, see Fig. 9A), the addition of nsp2^H114R^ rendered the rEAV^QYA^ virus nearly insensitive to ALV treatment, while also retaining its reduced CsA sensitivity (Fig. 10). These results clearly confirmed that the highest level of ALV resistance is achieved by introducing a combination of adaptive mutations in EAV nsp2 and nsp5.

**Fig. 10.**
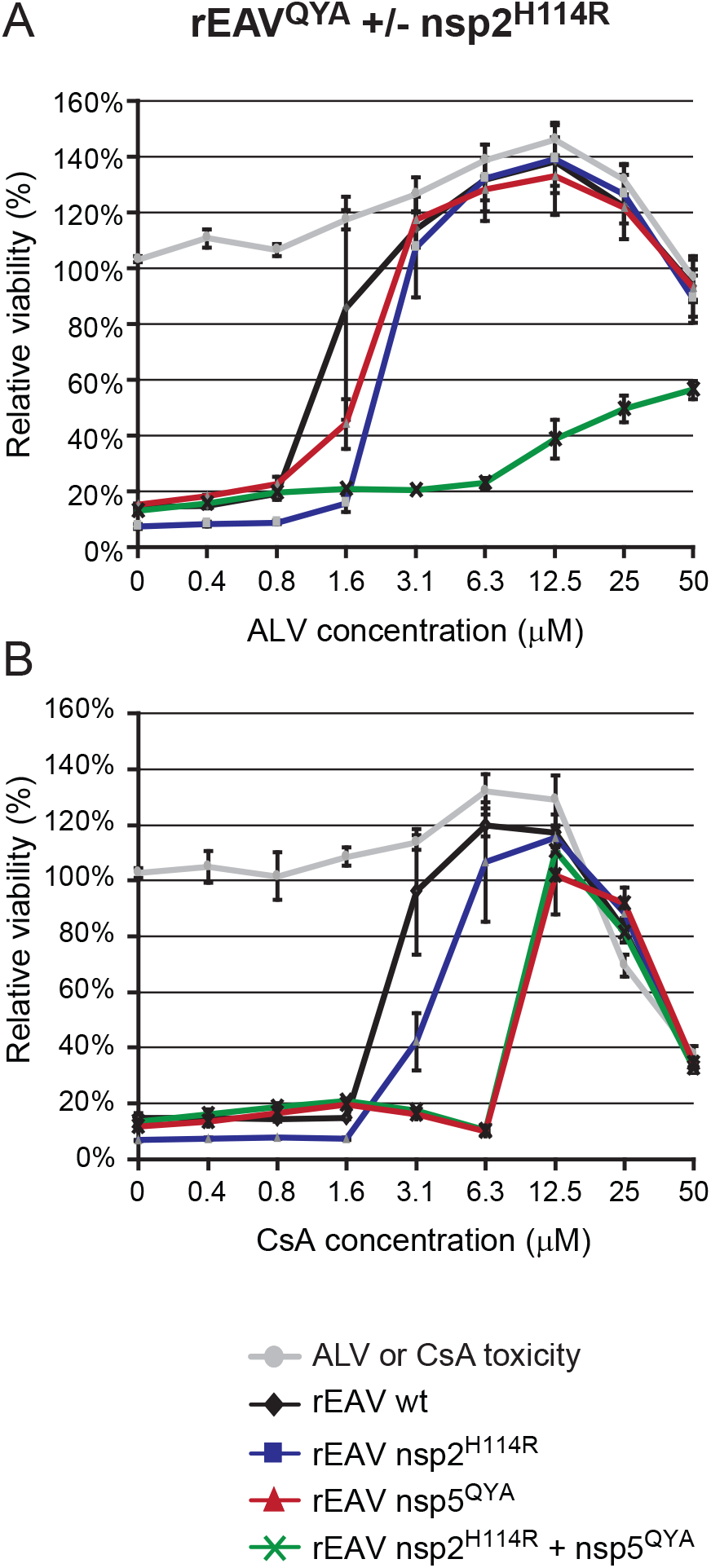
EAV resistance to ALV requires an additional mutation in nsp2. (A, B) Huh7 cells in 96- well plates were infected with rEAV^wt^ (control, black line) and the rEAV mutants rEAV-nsp2^H114R^ (blue), rEAV^QYA^ (red) and rEAV^QYA^-nsp2^H114R^ (green) in the presence of 0 to 50 µM ALV (A) or CsA (B). Cells were incubated for three days and cell viability was monitored using a commercial assay. In addition, the cytotoxicity of ALV or CsA treatment only was monitored in parallel in mock-infected Huh7 cells (light grey line). Graphs show the results (average and SD) of a representative experiment that was performed in quadruplicate. All experiments were repeated at least twice.

## Discussion

This study demonstrates that adaptive mutations in the transmembrane replicase subunit nsp5 can reduce the sensitivity of EAV replication to CsA treatment. Independently, experiments using the CsA analog ALV pointed in the same direction, in addition to implicating nsp2 in ALV sensitivity and resistance. Substitutions in nsp5 and nsp2 enable EAV to tolerate CsA or ALV concentrations that would normally block virus replication (Figs. 2, 9 and 10). The largest EC_50_ increase, about 5-fold, was achieved when multiple substitutions in nsp5 (and nsp2, in the case of ALV) were combined (Table 1 and 3). We previously showed that CsA inhibits EAV RNA synthesis and hypothesized that this may be the major determinant for its inhibitory effect (de Wilde et al., 2013a). Indeed, a combination of three of the acquired CsA resistance mutations in nsp5 (Q21R, Y113H, and A134V) facilitated a certain level of EAV RNA synthesis in the presence of a CsA dose that blocks wt virus activity. This was observed in both virus-infected cells and an *in vitro* assay for EAV RNA synthesis (Fig. 7).

Unfortunately, arterivirus nsp5 has not been studied in great detail and little is known about its functions and interactions during virus replication. The protein is one of three ORF1a-encoded transmembrane subunits that are presumed to drive the formation of the membranous ROs supporting viral RTC activity in the infected cell (Knoops et al., 2012; Pedersen et al., 1999; van der Hoeven et al., 2016). In expression systems, nsp2 and nsp3 interact and their combined expression suffices to induce the conversion of ER membranes into double-membrane vesicles that strikingly resemble those observed in arterivirus-infected cells. Although nsp5 appears to be dispensable for the basic interactions leading to these membrane transformations (Snijder et al., 2001), the protein was recently postulated to play a regulatory role by modulating membrane curvature and DMV formation (van der Hoeven et al., 2016).

The 162-amino acid nsp5 of EAV is largely hydrophobic and contains four predicted membrane-spanning domains and a cytosolic C-terminal domain (Fig. 1 and (van der Hoeven et al., 2016)). In infected cells, nsp5 is mainly present within a number of long-lived processing intermediates of the pp1a and pp1ab replicase polyproteins (Snijder et al., 1994; Wassenaar et al., 1997), in particular nsp5-7 and nsp3-8, but also larger intermediates. The latter extend into the ORF1b-encoded part of the replicase polyprotein, which includes key enzymes for arterivirus RNA synthesis (Snijder et al., 1994; Wassenaar et al., 1997). Consequently, nsp5 may have an important role as an ‘interaction platform’ that could target and/or anchor other RTC subunits to membranes. Being part of several larger processing intermediates, nsp5 may potentially perform such a role *in cis*, prior to the proteolytic release of those other subunits from nsp5-containing precursors. Obviously, also host cell proteins involved in arterivirus RNA synthesis are likely to become associated with the membrane-bound viral RTC, and may thus interact with nsp5. Such factors may include CypA and other members of the Cyp family that bind CsA and ALV, although the possibility of a direct interaction of these compounds with EAV replicase subunits themselves has not been formally excluded. In any case, our EM analysis of CsA-treated, EAV-infected cells, which were devoid of DMVs (Fig. 5), confirmed that the drug treatment blocks an early and basic step in the viral replicative cycle, and that this block can apparently be circumvented by the acquisition of a few adaptive mutations in nps5.

Four substitutions in nsp5 were implicated in CsA resistance (L8S, Q21R, Y113H, and A134V), of which only the L8 residue is conserved among arteriviruses (Fig. 1C). Interestingly, this residue is also part of a potential CypA binding motif (GPx**L**, as identified by (Piotukh et al., 2005)), constituting a potential interaction site between the two proteins. However, during infection, this L8S mutation quickly reverted, and therefore the contribution of the L8S substitution to CsA resistance remains unclear. All other three nsp5 mutations (Q21R, Y113H, and A134V) are present in predicted transmembrane domains (Fig. 1B), and therefore may not be directly involved in interactions of nsp5 with Cyps or other proteins. There is no experimental data available on the membrane topology of nsp5, but upon *in silico* modeling these nsp5 mutations did not appreciably alter the predicted conformation of the various transmembrane domains (data not shown). In addition, our EM analysis of DMV formation upon rEAV nsp5^QYA^ infection did not reveal any ultrastructural differences, either in the absence or presence of CsA (Fig 5). Clearly, more detailed information on the 3D-structure of nsp5 is needed to assess the impact of the mutations identified in this study on the protein’s conformation(s) and interactions at the molecular level.

We previously postulated a role for CypA in EAV RTC formation or function, upon demonstrating the cosedimentation of CypA with EAV RTCs in density gradients. Interestingly, no CypA cosedimentation with RTCs was detected in the presence of CsA (de Wilde et al., 2013a). Unfortunately, co-immunoprecipitation experiments using lysates from either ectopic expression of the EAV nsp2-7 polyprotein in 293T cells or EAV-infected cells, could not confirm an interaction between nsp5 and CsA, even when using mild conditions for cell lysis and immunoprecipitation (data not shown). This could obviously be due to technical reasons, e.g. the interaction being very weak or short-lived but an alternative explanation may be that the interaction of CypA with the EAV RTC is not directly via nsp5, or that it depends on factors that were lacking in our assays.

Although a direct interaction between an EAV nsp and CypA could not be established, efficient EAV replication is dependent on the presence of CypA, as it is severely reduced in cells depleted for CypA (Fig 6; (de Wilde et al., 2013a; de Wilde et al., 2018b)). For HCV, it was postulated that the interaction between CypA and NS5A is involved in the formation and stabilization of the RO, the so-called ‘membranous web’ with which viral RNA synthesis is associated (Chatterji et al., 2015; Madan et al., 2014). A direct interaction has been demonstrated between CypA and the proline-rich domain 2 of NS5A (Foster et al., 2011; Ngure et al., 2016). Presumably, CypA catalyzes the proper folding of NS5A, which in turn enhances the direct binding of RNA to NS5A, or recruitment of the NS5B-RdRp to the membrane structures, ultimately leading to efficient RNA synthesis (Coelmont et al., 2010). HCV mutations associated with resistance to Cyp inhibitors map to NS5A, close to the cleavage site between NS5A and NS5B, and either slow down processing of the NS5A-5B junction or render replication less dependent on CypA (Kaul et al., 2009). Interestingly, NS5A mutations that confer resistance to Cyp inhibitors do not reduce CypA-NS5A binding (Badillo et al., 2017). In analogy with these data, all relevant adaptive mutations in EAV nsp5 map inside the first and last of its four (predicted) transmembrane domains. One possible mechanism is that the adaptive nsp5 mutations somehow increase the stability of RTC complexes and thereby counteract the destabilization that may result from CsA penetrating lipid membranes. It has been shown that CsA can be inserted into and destabilize membrane structures, with a preference for sphingomyelin-rich membranes (Azouzi et al., 2010; Dynarowicz-Latka et al., 2015). In addition, several reports have been published that show that CsA treatment induces the unfolded protein response (UPR) and endoplasmic reticulum (ER) stress (Bouvier et al., 2009; Ram and Ramakrishna, 2014), and indeed, severe dilation of ER membranes was observed in CsA-treated Huh7 cells (compare for instance mock-infected cells in Figs. 4A and 4D). CsA may thus also affect the stability of virus-induced membrane structures and by doing so inhibit viral RNA synthesis, for example by disrupting the integrity of the viral RTC or its association with the ROs. Integration of mutant nsp5 in EAV RTCs may stabilize the RTC complex and allow it to tolerate higher CsA concentrations before it disassembles and loses its RNA synthesis activity.

Interestingly, despite the fact that CsA and ALV both are cyclophilin inhibitors, EAV resistance to ALV requires an additional mutation in the nsp2 protease domain PLP2: H114R (Fig 10). This suggests that, compared to CsA, ALV has an additional target that is relevant for EAV replication, either directly or indirectly. Residue H114 is positioned in helix α4 in domain I of the PLP2 structure, which is distant from both the protease’s active site and the surface involved in ubiquitin binding (van Kasteren et al., 2013), a feature linked to PLP2’s secondary function as a deubiquitinase involved in arteriviral innate immune evasion. Unfortunately, only the structure of the PLP2 domain of nsp2 has been resolved, and the overall fold and membrane topology of the subunit of EAV remains unclear. Our study implicates nsp5 in viral RNA synthesis, either directly or indirectly, and provides evidence that mutations in nsp5 are involved in reduced sensitivity to CsA, while the same mutations in combination with the nsp2 H114R substitution contribute to ALV resistance. Future studies should aim to elucidate the difference between the modes of action of ALV and CsA and the exact role of nsp5 and nsp2 in arterivirus sensitivity and resistance to Cyp inhibitors.

## Acknowledgements

We thank Ettore Ullo, Jayron Habibe, Jessika Zevenhoven-Dobbe and Diede Oudshoorn for technical assistance and helpful discussions, Dr. Grégoire Vuagniaux for critically reading the manuscript, and Novartis (Basel, Switzerland) and DebioPharm (Lausanne, Switzerland) for providing Alisporivir. This research was supported in part by and by the EU-FP7-Health project SILVER (grant 260644).

---

**sFig. 1.**
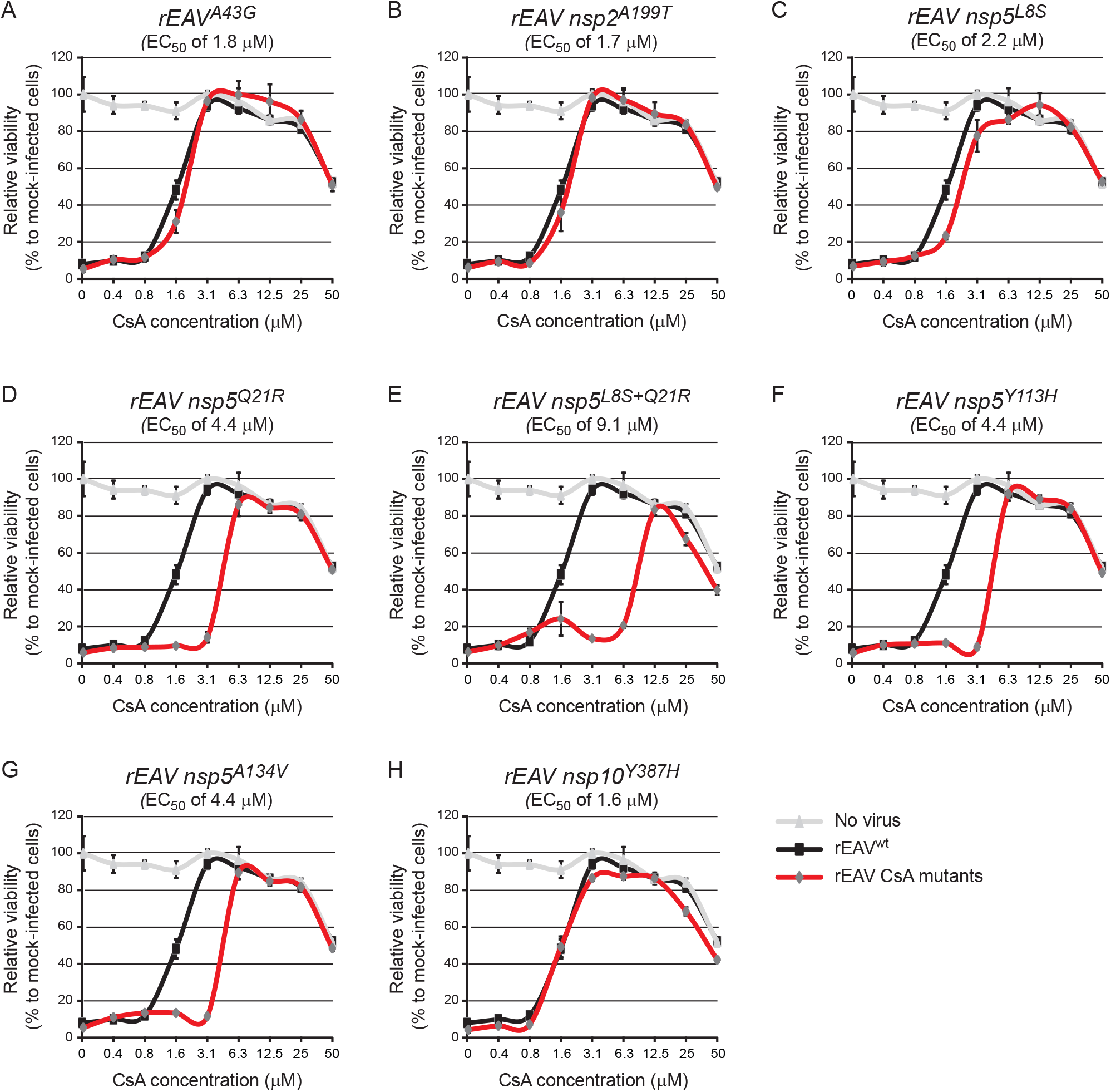
CsA-resistance analysis of rEAV mutants harboring singular mutations identified in the consensus sequences of CsA-1, -2, and -3. rEAV mutants with each of the individual mutations identified in the CsA-1, -2, and -3 consensus sequences were engineered and tested in a CPE-based assay. Huh7 cells in 96-well plates were infected (MOI 0.05) with rEAV^wt^ (control, black line) and rEAV CsA mutants (red line). Cells were incubated for three days in the presence of 0 to 50 µM CsA and cell viability was monitored using a commercial cell viability assay. The cytotoxicity of CsA treatment only was monitored in parallel in mock-infected Huh7 cells (light grey line). Above each panel, the EC_50_ value of CsA inhibition is indicated (see also Table 1). Graphs show the results (average and SD) of a representative experiment that was performed in quadruplicate. All experiments were repeated at least twice.

**sFig. 2.**
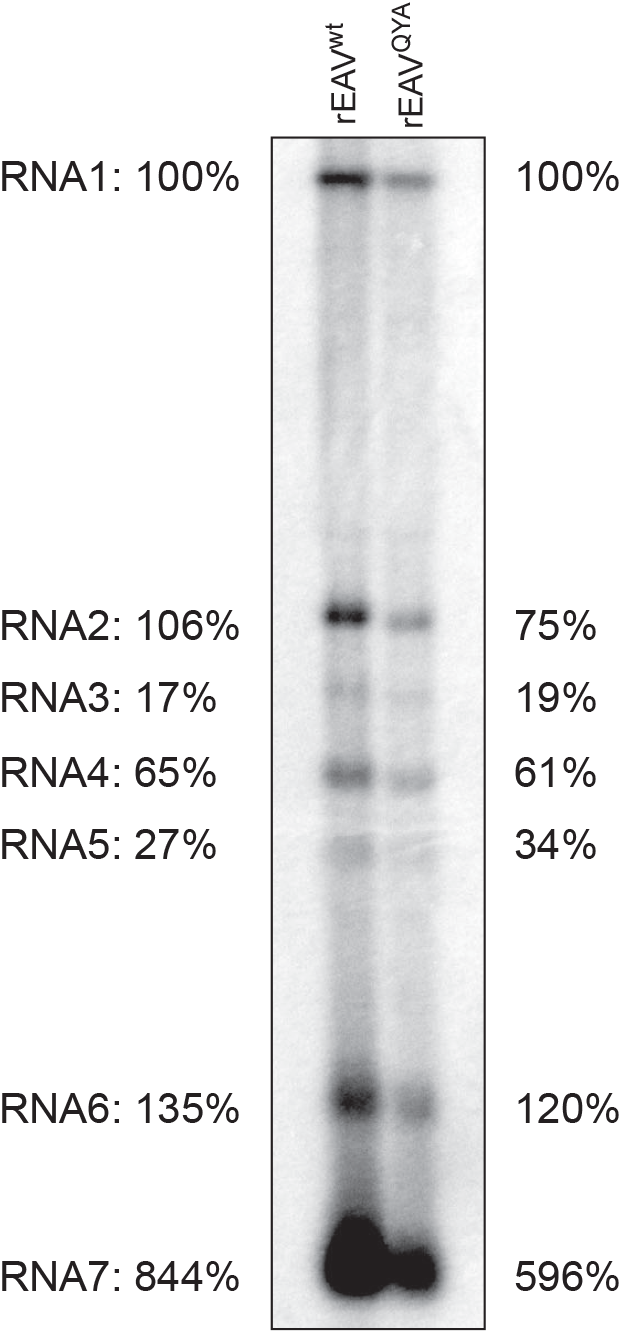
Hybridization analysis of RNA synthesis in rEAV^wt^- and rEAV^QYA^-infected cells. Intracellular RNA was isolated at 7.5 h p.i. from rEAV^wt^- and rEAV^QYA^-infected BHK-21 cells and analyzed in a denaturing formaldehyde agarose gel. The EAV RNA was visualized by hybridization to a ^32^P-labeled oligonucleotide probe (see Material and Methods) complementary to the 3’ end of EAV genome and sgRNAs. The positions of the genomic RNA (RNA1) and subgenomic RNAs 2 to 7 are indicated on the left side of the gel. Subgenomic RNA abundance was measured by phosphor imaging-based quantification of RNA bands and is given relative to the abundance of RNA1, which was placed at 100%.

